# Cell Type-Specific Changes in Dendritic Spines Across Adolescence Within Mouse Medial Prefrontal Cortex

**DOI:** 10.1101/2025.07.03.663087

**Authors:** Courtney M. Klappenbach, Qing Wang, Kylee Lenkersdorfer, Jacob Buursma, Jason “Jayes” Acuña, Jasmin “Ocean” Chu, Weihang Chen, Kaylie Richards, Ximena Herrera, Katy Touretsky, Kristen Delevich

## Abstract

Across species, cognitive capacities that rely on the frontal cortex do not fully mature until adulthood. Adolescent circuit refinement, including structural remodeling of dendritic spines, is believed to underlie this protracted maturation. Understanding cell type-dependent patterns of structural maturation would provide important insight into frontal cortex development. Here, we leveraged retrograde adeno-associated viruses to quantify dendritic spines on pyramidal tract (PT) vs. intratelencephalic (IT) neuronal populations in parallel within the mouse medial prefrontal cortex (mPFC) across adolescence. IT-type neurons showed opposing changes in mushroom and thin spines that were: 1) consistent with increasing synaptic maturity and 2) largely absent in PT-type neurons. We next probed the function of brain-resident immune cells, microglia, by transiently ablating them within the mPFC at mid-adolescence. This led to cell type-dependent changes in dendritic spines in late adolescence, with thin spine proportion increasing on both cell types but total spine density increasing on IT-type neurons only. Meanwhile, there was no effect on performance in an mPFC-dependent task of cognitive flexibility at either late adolescent or adult time points following microglia ablation. These findings provide evidence that mPFC IT-type neurons undergo greater spine remodeling during adolescence compared to PT-type neurons and implicate microglia as potential mediators.

## INTRODUCTION

The refinement of neural circuits within the frontal cortex during adolescence is believed to be critical for the emergence of mature cognitive function (Tervo-Clemmens et al. 2023; Piekarski et al. 2017; Klune, Jin, and DeNardo 2021; Larsen and Luna 2018; Chini and Hanganu-Opatz 2021). Within the rodent mPFC, glutamatergic excitatory pyramidal neurons play a pivotal role in integrating and transmitting information between cortical and subcortical areas (Anastasiades and Carter 2021). Two major subtypes of pyramidal neurons, IT-type and PT-type neurons, exhibit distinct morphology, electrophysiological properties, connectivity, and functional roles (Baker et al. 2018; Delevich, Thomas, and Wilbrecht 2018; Shepherd 2013; Moberg and Takahashi 2022). IT-type neurons primarily project within the cortex and to the striatum, facilitating interhemispheric communication, whereas PT-type neurons, also referred to as extratelencephalic (ET) neurons, project subcortically to regions like the brainstem and spinal cord, influencing motor output (Harris and Shepherd 2015; Cowan RL 1994; Reiner A and Del Mar N 2003; Naka A 2016; Baker et al. 2018). Within the mPFC, IT-type neurons can be found across Layers 2-6, intermingling with PT-type neurons within Layer 5 (Anastasiades and Carter 2021). Notably, while both IT- and PT-type neurons in the mPFC project to the striatum, only IT-type neurons cross the corpus callosum to innervate the contralateral striatum (Catsman-Berrevoets et al. 1980; Shepherd 2013; Morita, Im, and Kawaguchi 2019). The two pyramidal neuron classes exhibit asymmetric connectivity, with largely unidirectional inputs from IT-type to PT-type neurons (Kiritani et al. 2012; Brown and Hestrin 2009; Mieko Morishima and Kawaguchi 2006; M. Morishima et al. 2011). This circuit organization suggests that IT-type neurons exert top-down control over PT-type neuron output, ensuring that local computations shape descending motor commands to guide decision making.

Dendritic spines, which house the postsynaptic sites of most excitatory synapses, are key structures for synaptic plasticity and connectivity (Berry and Nedivi 2017; Bourne and Harris 2007). These micron-scale structures emanate from dendrites and contain the requisite molecular machinery for synaptic transmission and experience-dependent plasticity (Runge, Cardoso, and De Chevigny 2020). Adolescence, the transition from juvenile to adult, is marked by significant synaptic reorganization within the frontal cortices that is believed to be important for optimal function (Spear 2000; Paus, Keshavan, and Giedd 2008). Several previous studies have examined adolescent changes in mPFC dendritic spine density and dynamics on either IT (Pöpplau et al. 2023; Markham, Mullins, and Koenig 2013; Delevich, Okada, et al. 2020) or PT (Piekarski et al. 2017; Pattwell et al. 2016; Johnson CM, Peckler H, and Janak PH 2016) cell types, but it remains poorly understood how spine properties may differ between the two cell types across the adolescent transition. Furthermore, previous examination of IT-type neurons has been restricted to the upper cortical layers (Layer 2/3), so it remains unclear to what extent findings generalize across layers. Recent work in rodent models suggests that frontal cortical IT-type neurons may exhibit a more protracted maturation than PT-type neurons, perhaps reflecting their role in late-developing higher-order cognitive processing (Delevich, Thomas, and Wilbrecht 2018). Data from mouse frontal cortex suggests that synaptic inhibition onto L2/3 and 5 IT-type neurons is dynamic during the adolescent period between postnatal day (P)25 and 45, whereas inhibition onto L5 PT-type neurons remains stable (Vandenberg and Caporale 2015; Piekarski, Boivin, and Wilbrecht 2017). We hypothesized that IT-type neurons may also exhibit greater structural maturation during adolescence compared to PT-type neurons, reflecting their key role in the adolescent reorganization of the frontal cortex.

The central nervous system resident macrophages, microglia, have emerged as a potential sculptor of mPFC microcircuits during adolescence (Pöpplau et al. 2023; Chini and Hanganu-Opatz 2021; Mallya et al. 2019; Schalbetter et al. 2022). Microglia have been shown to participate in synaptic pruning (Paolicelli et al. 2011; Schafer et al. 2012), although the precise mechanisms by which they do so and their necessity in developmental synaptic refinement remains an active area of research (O’Keeffe et al. 2025; Pereira-Iglesias et al. 2025; Weinhard et al. 2018). There is evidence for increased microglia colocalization with dendritic spines during adolescence within the mPFC (Mallya et al. 2019; Pöpplau et al. 2023), and recent studies have shown that transient microglia ablation during adolescence, but not before or after, leads to long-term changes in pyramidal cell morphology and mPFC-dependent cognition (Schalbetter et al. 2022; Pöpplau et al. 2023). It is possible that microglia contribute to cell type and microcircuit differences in the timing and extent of structural remodeling, and we therefore sought to examine the role of microglia in shaping structural maturation of mPFC IT vs. PT-type neurons.

In the current study, we quantified dendritic spines on projection-defined PT-type vs. IT-type neuronal populations in parallel within the mPFC across adolescence. Our goal was to examine whether PT-type and IT-type neurons exhibit different developmental trajectories, with the hypothesis that IT-type neurons may exhibit greater dendritic spine changes during adolescence compared to PT-type neurons. Next, we examined whether focal ablation of microglia within the prelimbic region of the mPFC during mid-adolescence would have cell type-dependent effects on spine properties and impact cognitive flexibility during late adolescence and adulthood.

## 1: METHODS

### 1.1: Animals

Male and female C57BL/6 mice (Charles River) were bred in-house. All mice were weaned on postnatal day (P)21 and housed in groups of 2–3 same-sex siblings on a 12:12 h reversed light:dark cycle (lights on at 1900 h) in standard ventilated polycarbonate cages with Biofresh cellulose bedding. Mice were provided ad libitum access to standard chow (Purina 5001) that contains ∼5% fat and water in the home cage. All procedures were approved by the Washington State University Institutional Animal Care and Use Committee and conformed to principles outlined by the NIH Guide for the Care and Use of Laboratory Animals. Full details of animal numbers for each experiment are provided in Supplementary Material. See Supp. Fig. 1 for experimental overview.

### 1.2: Stereotaxic Injections

Male and female mice were deeply anesthetized with 5% isoflurane (vol/vol) in oxygen and placed into a stereotactic frame (Kopf Instruments; Tujunga, CA) upon a heating pad. Anesthesia was maintained at 1%–2% isoflurane during surgery. An incision was made along the midline of the scalp and small burr holes were drilled over each injection site. Virus was delivered via microinjection using a Nanoject II injector (Drummond Scientific Company; Broomall, PA). As previously described, retrograde AAV was delivered into the dorsomedial striatum (DMS) and pons to label IT-type and PT-type neurons, respectively (Delevich, Okada, et al. 2020). Specifically, 50 nL of retrograde AAV virus (pAAV-CAG-GFP [Addgene item #37825] or pAAV-CAG-tdTomato [ Addgene item #59462]) diluted 1:5 (GFP) or 1:10 (tdTomato) in saline (starting titer ≥7×10^12^ vg/mL) was delivered unilaterally to the DMS (coordinates relative to Bregma: A/P: +0.9 mm, M/L: +1.4 mm, D/V: +3.0 mm for adults, +2.8 mm for juveniles) and pons (coordinates relative to Bregma: A/P: −4.26 mm, M/L: +0.6 mm, D/V: -4.6 mm for adults, -4.3 for juveniles). Mice were given subcutaneous injections of meloxicam (0.05 mg/kg) and buprenorphine HCl (5mg/kg) prior to surgery as well as 24 and 48 h after surgery. Mice were group housed before and after surgery. For cell mapping and spine imaging in Experiments 1 & 2, fluorophore and hemisphere were counterbalanced across mice, with DMS and pons injections always performed in opposite hemispheres. For Experiment 3, 200 nl of 50 mg/ml clodronate disodium salt (CDS, MedChemExpress, #HY-B0657A) or Dulbecco’s phosphate-buffered saline (DPBS, Corning, #21-030-CV) was injected into the prelimbic region (coordinates relative to Bregma: A/P = +2.1 mm, M/L = +0.4 mm, D/V = -1.5 mm) unilaterally at P44 to quantify microglia ablation. For Experiment 4, CDS or DPBS were injected ipsilateral to the pons injection and contralateral to the DMS injection in mice at P44. For Experiment 5, CDS or DPBS were injected bilaterally into the prelimbic region at P44.

### 1.3: Histology

Eight days after virus injection, mice were transcardially perfused with 4% paraformaldehyde (PFA) in 0.1 M phosphate buffer (PB) solution (pH=7.4). Brains were harvested and postfixed in 4% PFA for 4 hours, followed by transfer to 25% sucrose in PB. For cell distribution and colocalization experiments, sections were cut to 50 µm using a cryostat (Leica, Deer Park, IL). For spine imaging, sections were cut at 100 µm. For both experiments, sections were blocked (0.2% TritonX-100, 5% normal goat serum [NGS, Jackson #005-000-121], 3% bovine serum albumin [BSA, Millipore #126579] in PB) for 2 hours at room temperature while shaking then incubated with a GFP booster (Chromotek, gba488) at a 1:200 dilution overnight in blocking solution (5% NGS, 2% BSA in PB). Sections were rinsed 4x with PB for 20 min each then mounted in Prolong Gold.

To verify and map DMS and pons injection sites, coronal sections containing these regions were imaged on a Zeiss Axio Imager M2 microscope with a 2.5x objective or a Leica Dmi8 microscope with a 5x objective. Images were imported into Adobe Illustrator and mapped onto the Paxinos mouse brain atlas (Paxinos and Franklin 2019) (see Supp. Figs. 2-3).

To verify the effect of CDS on microglia, mice injected with CDS or DPBS alone were transcardially perfused after five days, a time point that was previously shown to coincide with maximal depletion of microglia following intracranial injection (Schalbetter et al. 2022). Sections were cut at 100 µm and stored in anti-freeze (50% 0.05M PB, 30% ethylene glycol, 20% glycerol at pH 7.3) at - 20°C until all sections had been collected so that they could be stained concurrently. Sections were rinsed 2x with PB for 10 min each then stained using metal enhanced DAB substrate kit (Thermo Fisher, SKU# 34065) and ABC peroxidase kit (Thermo Fisher, SKU# 32020) following kit protocol. All incubations were done at room temperature while shaking. Briefly, sections were incubated with a blocking buffer (with TritonX-100, described above) for 2 h then incubated with a 1:3000 dilution of rabbit anti ionized calcium-binding adaptor molecule 1 (Iba-1) primary antibody (Fuji Film, 018-28523) in blocking solution (without TritonX-100, described above) overnight. Sections were then rinsed 3x with PB for 5 min each before being incubated with a 1:500 dilution of goat anti rabbit biotin conjugated secondary antibody (Thermo Fisher, A16100) overnight. Sections were again washed 3x with PB for 5 minutes each then incubated with the ExtrAvidin-peroxidase (XAV, Millipore-Sigma, #E2886) solution at a 1:1500 dilution for 4 hours. Sections were washed again then incubated with the filtered DNB/Nickel solution (25% 0.4M PB [pH 7.4], 1% NH_4_Cl [Sigma, #A9434], 0.2% glucose, 0.5 mg/ml 3,3′-Diaminobenzidine [DAB, Millipore Sigma, #D8001], 0.04% ammonium nickel (II) sulfate hexahydrate [Sigma, #A1827]) for 10 minutes then 1 µl of glucose oxidase (Millipore-Sigma, #A1827) was added for 8 min before quenching the reaction by rinsing 3x with PB. Samples were mounted in Permount (Fisher Scientific, #SP15-100) overnight with dehydrated coverslips. Coverslips were dehydrated by moving between the following solutions subsequently for 2 min each: 95% EtOH x2, 100% EtOH x2, 2-propanol x2, SafeClear x2 [Protocol, #314-629].

### 1.4: Fluorescence Microscopy for Cell Distributions

For comparisons of IT-type and PT-type neuron distribution, coronal sections were imaged on a Zeiss Axio Imager M2 microscope with a 5x objective or a Leica Dmi8 microscope with a 20x objective. Three sections containing mPFC were imaged per mouse. Cell counts and colocalization of retrogradely-labeled neurons were performed in ImageJ using the ‘Cell Counter’ plugin with different counter types for red (tdTomato-labeled), green (GFP-labeled), and yellow (co-labeled) cells. Cells were counted only if they were located within the prelimbic and anterior cingulate portions of the mPFC. The X and Y position of each neuron was exported to CSV files in addition to the X coordinate of the midline of the coronal section. This data was processed via an in-house R script to output the distance of every neuron to the midline and reference point (defined as the intersection of the midline and the most dorsal point of the brain section).

### 1.5: Confocal Imaging for Dendritic Spines

Dendritic spines were imaged on a Leica SP8X confocal microscope with LIGHTNING deconvolution and a 63x oil immersion lens. All images were taken with frame sequential scanning, a step size of 0.2 µm, and approximately 60 - 80 steps. The 488 nm laser was set to 7% power and the 554 nm laser was set to 10% power. Images were taken in the hemisphere that was contralateral to the DMS injection and ipsilateral to the pons injection. Images were taken so that the section midline (medial wall of mPFC) was visible at the edge of the field of view. All images were 144.78 µm^2^, which ensured that all dendrites imaged and scored were within 144.78 µm of the pia surface. Two images were taken per section comprising both red and green channels, and three sections were imaged per mouse. This resulted in 12 total images per mouse (3 sections x 2 images x 2 fluorophores). One mouse (P29 male) had incorrect virus targeting for the DMS injection site and therefore only PT-type dendrites were imaged and scored.

To determine the exact location of each confocal image within the mPFC, a bleach spot was created in a distal location to each region of interest (ROI) by scanning the 405 nm laser at 100% power for 1-2 minutes. The coordinates of each ROI and bleach spot were recorded and used to map the imaging locations to the Paxinos brain atlas using low magnification overview images of the sections (see Supp. Fig 4).

The first analysis of spine parameters performed was spine density, presented as spines per 10 µm of dendrite. Our initial statistical model included fixed effects of sex, age, cell type, and fluorophore. The model also included nested random effects to account for the repeated measures within each mouse, brain section, and image / field of view. While fluorophore color (tdTomato vs. GFP) was counterbalanced across groups, we included it in the model to determine if it accounted for additional variation. Unexpectedly, we discovered a significant main effect of fluorophore (*p* < .0001) where GFP labeled dendrites were scored as having significantly higher spine density than tdTomato labeled dendrites (GFP labeled = 3.87 ± .0944, tdTomato labeled = 3.34 ± .0733). This was true across both cell types and all ages (Supp. Fig. 5A-B). The only interaction detected with fluorophore was sex (*p* = .0345), where the magnitude of the fluorophore discrepancy was larger in female than male mice, however GFP labeled dendrites were still scored as denser than tdTomato labeled dendrites (Supp. Fig. 5C). Due to limited significant interactions with other main effects and consistent patterns across age and cell type, we included fluorophore as an additive / covariate factor in all subsequent models. We do not further discuss fluorophore as a significant main effect, however, this information can be found in the Supplementary Statistics file.

### 1.6: Brightfield imaging and stereology of DAB-stained IBA-1 sections

All imaging and stereology were performed using a Zeiss Axio Imager M2 microscope with StereoInvestigator version 2022.3.1. Representative section images were taken using a 2.5x or 5x objective. Although only one hemisphere was injected with CDS or DBPS, both hemispheres were scored because of observed interhemispheric extent of microglia ablation.

### 1.7: Spine Image Processing / Analysis

Images were blinded and imported into Neurolucida 360 (MBF Biosciences) version 2018.1.1. Some images were excluded from scoring either due to a lack of dendrites or because the density of labeled dendrites was too high. Two dendrites for each channel (tdTomato or GFP) per image were traced manually using ‘Smart Manual’ mode and the ‘Directional Kernels’ method. All dendrites were traced by the same experimenter. Spines were scored using the click to detect method. Spine type was determined using the ‘Classify all’ function in Neurolucida 360 and default classification parameters (Rodriguez et al. 2008). Images were distributed to three experimenters for scoring in sets of 50-100. A subset of images was scored twice within each assigned image set to estimate intra-set, intra-experimenter variability. A second subset of images was then scored to estimate inter-set intra-experimenter variability. Finally, a third subset of images were scored by multiple experimenters to estimate inter-experimenter variability. The average difference ± standard deviation in spine density (spines / µm) was as follows: intra-set intra-experimenter = 0.062 ± 0.052 (n = 60 dendrites), inter-set intra-experimenter = 0.072 ± 0.062 (n = 55 dendrites), inter-experimenter = 0.067 ± 0.055 (n = 32 dendrites). A batch analysis of scored images was performed in Neurolucida Explorer version 2018.2.1 using the Dendritic Segments, Dendritic Spines, and Spine Details analysis options. The output excel files from this analysis were compiled, unblinded, and analyzed in R.

### 1.8: Cognitive flexibility testing following adolescent microglia ablation

Male and female C57Bl/6 mice were bilaterally transfused with CDS or DPBS into the prelimbic region at P44. Next, after 8 days (P52) or 6 weeks (P86), mice began training in the four-choice odor based task, as previously described (Delevich et al. 2022; Delevich, Hall, et al. 2020; C. Johnson and Wilbrecht 2011). Briefly, mice were food restricted to either 95-100% body weight (late adolescent, P52) or ∼85-90% body weight (adult, P86) by the discrimination phase. All tests were conducted under red light by an experimenter blinded to treatment. On day 1, mice were habituated to the testing arena (habituation), on day 2 were taught to dig for a 20 mg grain pellet (TestDiet 5TUM) reward in a pot filled with unscented wood shavings (shaping), on day 3 underwent a four-choice odor discrimination, and finally on day 4 were tested on recall of the previously learned odor discrimination immediately followed by a reversal phase. During the discrimination phase of the task, mice learned to discriminate among four pots with different scented wood shavings (anise, clove, litsea, and thyme). All four pots were sham-baited with grain pellets (under wire mesh at the bottom) but only one pot was rewarded (anise). The pots of scented shavings were placed in each corner of an acrylic arena (12″ L, 12″ W, 9″ H) divided into four quadrants. Mice were placed in a cylinder in the center of the arena, and a trial started when the cylinder was lifted. Mice were then free to explore the arena and indicate their choice by making a bi-manual dig in one of the four pots of wood shavings. The cylinder was lowered as soon as a choice was made. If the choice was incorrect, the trial was terminated and the mouse was gently encouraged back into the start cylinder. Trials in which no choice was made within 3 min were considered omissions. If mice omitted for two consecutive trials, they received a reminder: a baited pot of unscented wood shavings was placed in the center cylinder and mice dug for the “free” reward. Mice were disqualified if they committed four pairs of omissions. The location of the four odor scented pots was shuffled on each trial, and criterion was met when the mouse completed 8 out of 10 consecutive trials correctly. Twenty-four hours after completing discrimination, mice were tested for recall of the initial odor discrimination to criterion. Immediately after reaching criterion in the recall phase, mice proceeded to the reversal phase in which the previously rewarded odor (anise) was no longer rewarded, a previously unrewarded odor (clove) now became rewarded, and another previous odor (thyme) was replaced by a novel odor (eucalyptus) that was unrewarded. Again, mice were run until they reached a criterion of 8 out of 10 consecutive correct trials.

### 1.9: Statistics

Statistical analysis was performed using linear mixed effects models in R using the lme4 package. Significance tests of model effects and interactions were performed with the emmeans package. Model effects and interactions were calculated with the joint_tests() function and *post hoc* comparisons were done with the emmeans() function and specifying pairwise comparisons. Group comparisons were combined with the rbind() function and the adjustment method ‘none’ was applied because they were restricted to a small set of planned comparisons. Models were specified with a hierarchical nested random effect of mouse, brain section, and image / field of view. For analysis of proportion of each spine type, the proportion was calculated across each brain section and therefore the image / field of view level of random effect was not included. Unless otherwise specified, values presented represent mean ± SEM. See supplemental statistics file for full details.

## 2: RESULTS

### 2.1: Spatial distribution of retrogradely-labeled IT- and PT-type neurons within mPFC

Mice received dual infusions of AAVretro GFP and tdTomato into the DMS (Fig. 1A) and pons (Fig. 1B). Given that only the axons of IT-type neurons project to the contralateral hemisphere of DMS, this labeling strategy resulted in IT_DMS_ and PT_pons_ soma labeling in the medial prefrontal cortex (mPFC) hemisphere contralateral to the DMS injection site (Fig. 1C). We previously demonstrated that this strategy labeled cell populations that exhibited membrane properties characteristic of IT- and PT-type neurons (Delevich, Okada, et al. 2020). IT_DMS_ and PT_pons_ neurons rarely colocalized regardless of hemisphere, indicating that axons of mPFC pyramidal neurons rarely collateralize to pons and DMS. IT_DMS_ neurons were significantly more superficial compared to PT_pons_ neurons (Fig. 1D-E), with a significant main effect of cell type on distance from midline (*p* < .0001) but not hemisphere relative to injection site (*p* = .23) or interaction between cell type and hemisphere (*p* = .75). These results suggest that the laminar position of callosally-projecting IT_DMS_ neurons was not significantly shifted relative to ipsilateral IT_DMS_ neurons.

**Fig. 1:**
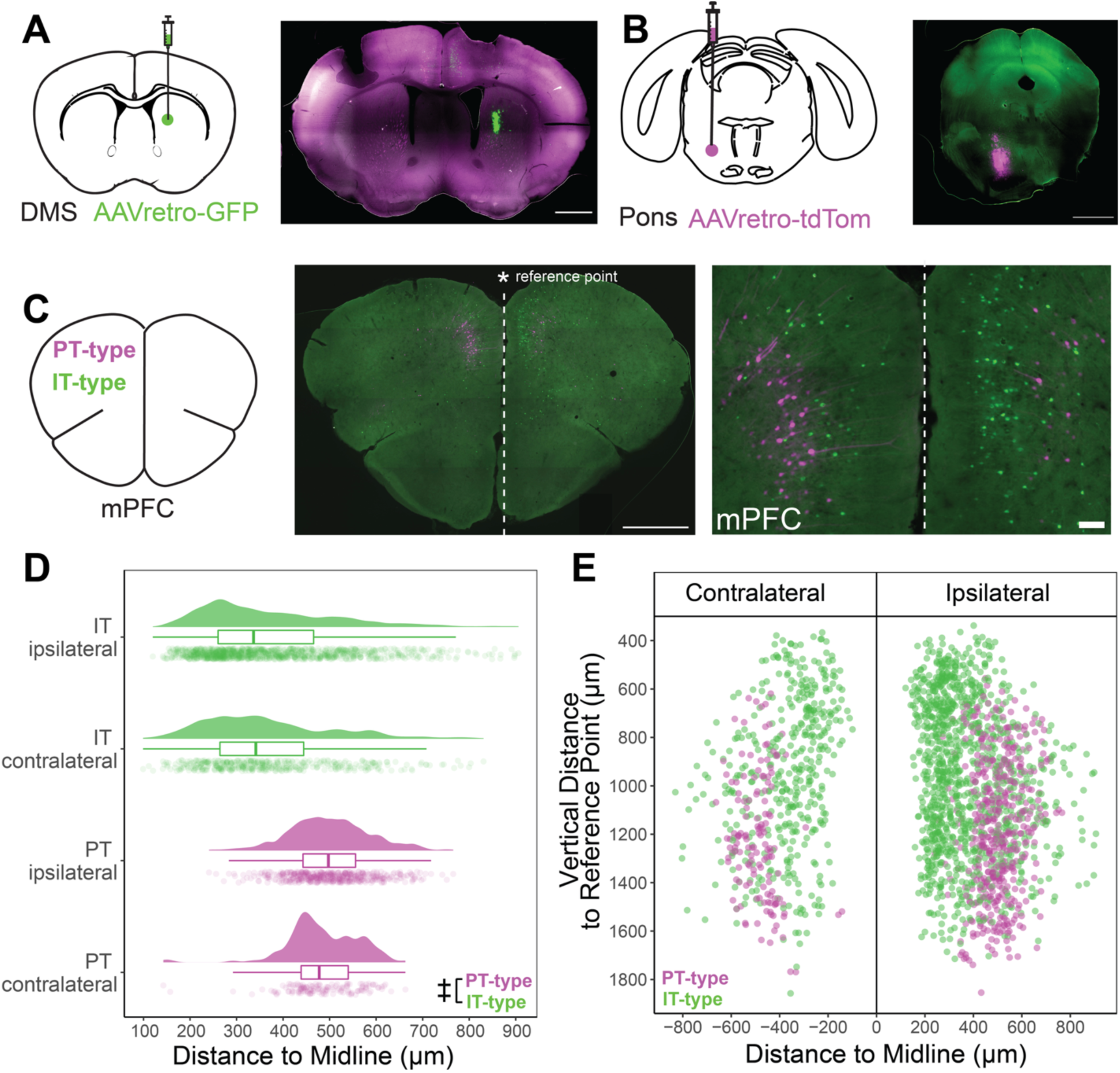
Dual AAVretro injections to DMS and pons label non-overlapping populations of pyramidal neurons within the mPFC. **A:** AAVretro virus of one fluorophore (GFP depicted) was injected into the DMS to label IT cells. **B:** AAVretro virus of the other fluorophore (tdTomato depicted) was injected into the pons to label PT cells (in the opposite hemisphere of the DMS injection). **C:** Representative image of retrogradely-labeled neurons in mPFC. Dashed line represents the midline and asterisk indicates the ‘reference point’. **D:** Distance to the midline for both cell types and relationship of neuron soma to injection hemisphere (ipsilateral or contralateral). ‡ indicates *p* < .0001 from linear mixed model with main effects and interactions between cell type and hemisphere. **F:** Spatial plot showing position of all somas as their distance to the reference point vs. distance to the midline. n = 4 mice, 12 sections, and 2093 neurons. Scale bars in whole section overviews indicate 1000 µm, scale bar for panel C inset indicates 100 µm.

### 2.2: Cell type-specific changes in spine density, spine type, and morphology across adolescence

To assess changes in spine parameters on IT_DMS_ vs. PT_pons_ neurons across adolescence, we performed dual AAVretro infusions into pons and DMS at postnatal day (P)21, P36, or P52, and collected brains 8 days later at P29, P44, and P60 (**Fig. 2**). These ages corresponded to late juvenile/early adolescent, mid-adolescent, and young adult time points, respectively (Brust, Schindler, and Lewejohann 2015). Confocal images of apical dendrites were collected from the mPFC hemisphere ipsilateral to the pons and contralateral to the DMS injection sites to compare callosally-projecting IT_DMS_ and PT_pons_ neurons within the same fields of view (Fig. 2A–D). There was a significant main effect of cell type (*p* < .0001) on apical spine density, with IT_DMS_ neurons having significantly higher spine density than PT_pons_ neurons (Fig. 2E). In addition, there was a significant interaction between cell type and sex (*p* = .02), whereby the cell type difference in spine density (IT_DMS_ > PT_pons_) was greater in females than males (Supp. 6A). Despite this interaction, *post hoc* sex comparisons within cell type were not significant (*ps* > .05). While the main effect of age was not significant (*p* = .11), there was a trend in both cell types where spine density increased from the P29 to P44 age group, then decreased from the P44 to P60 (Fig. 2E).

**Fig. 2:**
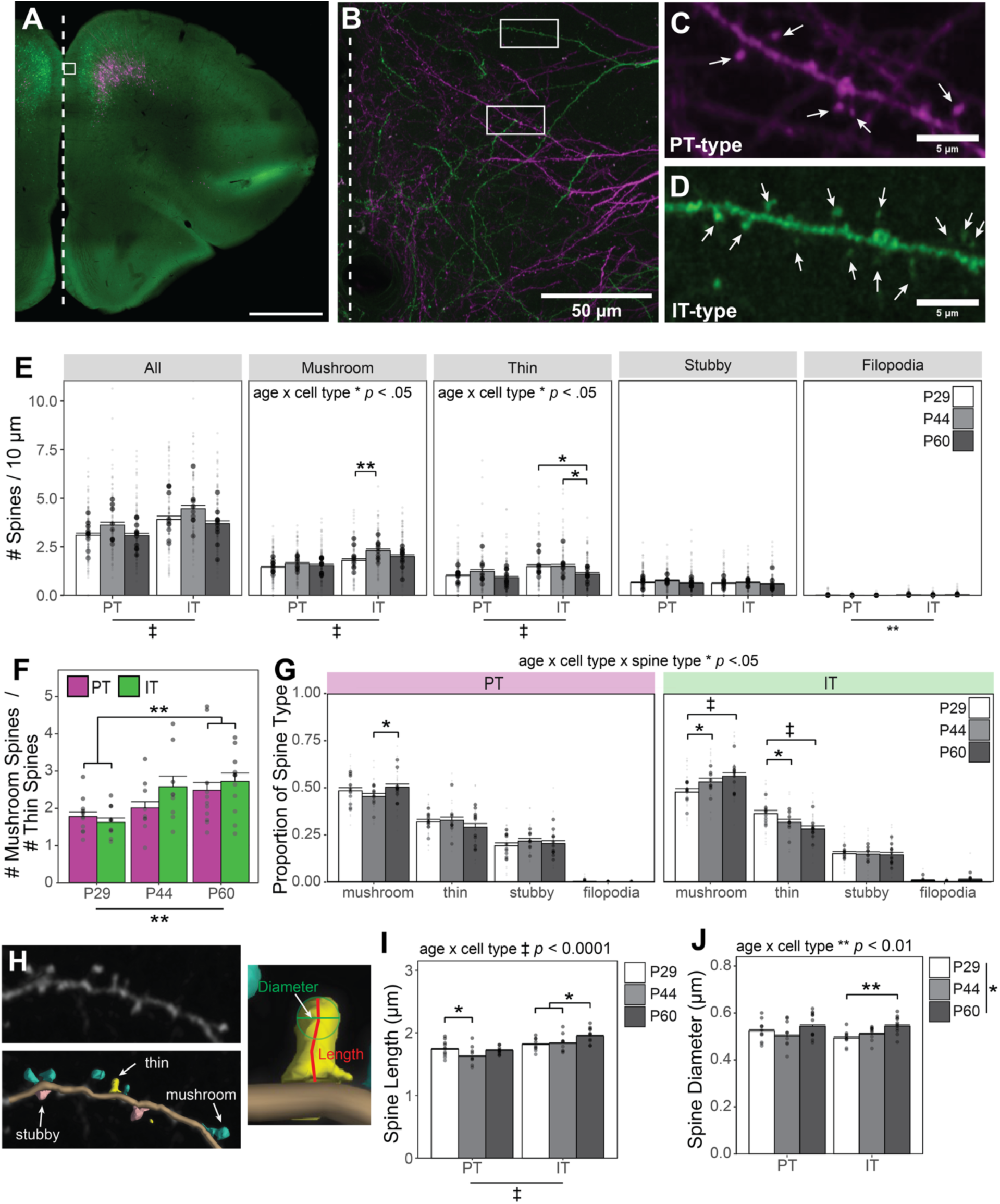
Cell type-specific changes in spine density and morphology across adolescence. **A:** Representative overview of dual AAVretro injected brain section including mPFC. Box shows size and representative location of confocal images. Scale bar indicates 1000 µm. **B:** Representative confocal image of apical dendrites. In both A and B, dashed line indicates midline. Boxes in B represent expanded dendrites from **C:** PT-type and **D:** IT-type neurons. Spines are denoted with arrows. **E:** Spine density for all spine types and for each class of spine type. **F:** Ratio of mushroom to thin spines. **G:** Proportion of each spine type was calculated across all dendrites within a brain section, so that the proportions of each spine type for a single brain section add to 1. **H:** Representative dendrite from Neurolucida 360 (NL360) with and without traced dendrite and spines. Labels and arrows indicate spine types as classified by NL360. Expanded image of spine shows backbone length and head diameter. **I:** Spine backbone length. **J:** Spine head diameter. Bar and error bars represent mean ± SEM of spine parameters for all spines. Small points represent individual dendrites and larger points represent means for each mouse (including F, I, and J). n = 9-11 mice / group, 27-33 sections / group, >100 dendrites / group, >1500 spines / group. For all panels, * indicates *p* < .05, ** indicates *p* < .01, *** indicates *p* < .001, and ‡ indicates *p* < .0001 for main effects, interaction effects, and *post hoc* comparisons from linear mixed effects models.

We next examined spine density by morphological class. For mushroom spines, there was a significant main effect of cell type (*p* < .0001) and a significant interaction between cell type and age (*p* < .0275) (Fig. 2E). *Post hoc* comparisons showed that the density of mushroom spines was higher on IT_DMS_ neurons compared to PT_pons_ neurons at all ages (*p*s < .01), and only IT_DMS_ neurons showed a significant effect of age on mushroom spine density (*p* = .0048). Thin spine density also showed a significant main effect of cell type (*p* < .0001) and interaction between cell type and age (*p* = .0172). Thin spine density was significantly higher on IT_DMS_ compared to PT_pons_ neurons at P29 (*p* < .0001) and P44 (*p* = .0103) but not P60 (*p* = .248). *Post hoc* comparisons showed that IT_DMS_ but not PT_pons_ cells had significantly lower thin spine density at P60 compared to P29 (*p* = .0202) and P44 (*p* = .0222). Stubby spine density showed no significant main effects or interactions. The model for filopodia spine density showed a significant main effect of cell type only (*p* = .0042), where IT_DMS_ neurons had a higher density of filopodia spines than PT_pons_ neurons, however both densities were extremely low (Fig. 2E). Collectively, spine density analyses revealed that IT_DMS_ neurons have a higher spine density than PT_pons_ cells which was true for all morphological classes except stubby spines. In addition, these results indicate that the density of mushroom and thin spines significantly changes across adolescence on IT_DMS_ but not PT_pons_ neurons.

Given the significant age effects on mushroom and thin spine density, we next examined the ratio of mushroom to thin spines, which is an indicator of spine maturation. The linear mixed effects model showed a significant main effect of age (*p* = .0094), with the ratio of mushroom to thin spines increasing with age (Fig. 2F). *Post hoc* comparisons showed that the mushroom/thin ratio was significantly higher at P60 compared to P29 in both cell types (*p* = .0025).

To further investigate how spine properties change with age, we compared the proportion of each morphological class (aka spine type) at P29, P44, and P60 (Fig. 2G). There was a significant 3way interaction among age, cell type, and spine type (*p* = .0262) as well as a significant 2way interaction between spine type and sex (*p* < .0001). To investigate the sex x spine type interaction, *post hoc* comparisons were run between males and females for each spine type (Supp. Fig. 6B). These showed that males had a higher proportion of mushroom spines than females (*p* = .0019) and that females had a higher proportion of thin spines compared to males (*p* = .0003).

To investigate the age x cell type x spine type interaction, *post hoc* comparisons between age groups within each cell type and spine type were performed. For PT_pons_ cells, only one *post hoc* comparison was significant (*p* = .0127), indicating that PT_pons_ cells do not drastically change the proportion of each spine type with age. For IT_DMS_ neurons, however, the proportion of mushroom spines significantly increased with age, with P29 being lower compared to P44 (*p* = .0157) and P60 (*p* < .0001). Conversely, the proportion of thin spines on IT_DMS_ neurons decreased with age, with the P29 group being significantly higher than P44 (*p* = .0428) and P60 groups (*p* < .0001). Neither cell type showed significant age-related changes in the proportion of stubby or filopodia spines.

We also performed *post hoc* tests to compare IT_DMS_ and PT_pons_ spine type proportions within each age group. At P44 and P60, IT_DMS_ neurons had a higher proportion of mushroom spines than PT_pons_ neurons (P44: *p* = .0003; P60: *p* = .0021). IT_DMS_ neurons had a lower proportion of stubby spines than PT_pons_ neurons at all ages (P29: *p* = .0387; P44: *p* = .0017; P60: *p* = .0022). Finally, at P29 only, IT_DMS_ dendrites had a higher proportion of thin spines compared to PT_pons_ dendrites (*p* = .0392). These results show that at P44 and beyond, IT_DMS_ dendrites have a higher proportion of mature (mushroom) spines and a lower proportion maturing spines (stubby) compared to PT_pons_ neurons.

To further assess underlying changes in spine type, we examined spine backbone length and spine head diameter (Fig. 2H). Analysis of spine backbone length (Fig. 2I) showed a significant main effect of cell type and 2way interactions of cell type x sex and cell type x age (*p*s <.0001). To explore the interaction of cell type with age, we performed *post hoc* comparisons between cell types within each age and between ages within both cell types. At all ages, spines on IT_DMS_ neurons had a longer backbone length than spines on PT_pons_ neurons (P29: *p* = .0008; P44: *p* < .0001; P60: *p* < .0001). Spines on PT_pons_ neurons showed a U-shaped pattern over time, getting shorter from P29 to P44, then longer again at P60 (P29 vs P44: *p* = .0276, P44 vs P60: *p* = .0822). IT_DMS_ cell spines showed a strong increase in backbone length with age (P29 vs P60: *p* =.0057, P44 vs P60: *p* = .0206). Exploration of the interaction between sex and cell type (Supp. Fig. 6C) showed that in both males and females, spines from IT_DMS_ cells again had longer backbone length than spines on PT_pons_ cells (females: *p* < .0001; males: *p* < .0001). Although males tended to have longer spines on IT_DMS_ cells than females, this difference was not significant (*p* = .06).

Analysis of spine head diameter (Fig. 2J) showed a significant main effect of age (*p* = .0392) and a significant 2way interaction between age and cell type (*p* = .0025). PT_pons_ spines again showed a U-shaped pattern, however no *post hoc* comparisons were significant. Spines on IT_DMS_ neurons showed an increase in spine head diameter with age, with IT_DMS_ spine heads significantly larger at P60 compared to P29 (*p* = .0024). Comparisons of cell type within each age showed that spines on PT_pons_ cells were longer than those on IT_DMS_ neurons only at P29 (*p* = .0007). Analysis of morphological parameters revealed that spines on IT_DMS_ neurons were longer compared to PT_pons_ spine and increased in length and diameter with age, whereas spine morphology on PT_pons_ neurons was relatively stable across adolescence.

### 2.3: Role of microglia in cell type-specific spine pruning

Recent data suggest that the brain resident immune cells, microglia, play a role in synaptic refinement within the mPFC during adolescence. We tested whether focal ablation of microglia within the mPFC at mid-adolescence (P44) would affect dendritic spines on IT_DMS_ and PT_pons_ neurons when measured in young adulthood. An initial cohort of mice was perfused 5 days following unilateral infusion of 200 nL chlodronate disodium salt (CDS) or DPBS vehicle (Fig. 3A-B). Consistent with a previous report (Schalbetter et al. 2022), we observed a significant reduction in the density of Iba-1+ microglia within mPFC of CDS injected mice compared to DPBS vehicle injected mice (*p* < 0.001) (Fig. 3C). A second cohort of mice received dual retroAAV injections into the DMS and pons as before at P44 in addition to a unilateral DPBS or CDS into the mPFC (Fig. 3D). Mice were perfused and brains collected 8 days later and dendritic spines on IT_DMS_ and PT_pons_ neurons were imaged (See Supp. Fig. 1D for timeline). Based on previous studies, we expected that microglia density would largely recover by 8 days post-injection (Schalbetter et al. 2022).

**Fig. 3:**
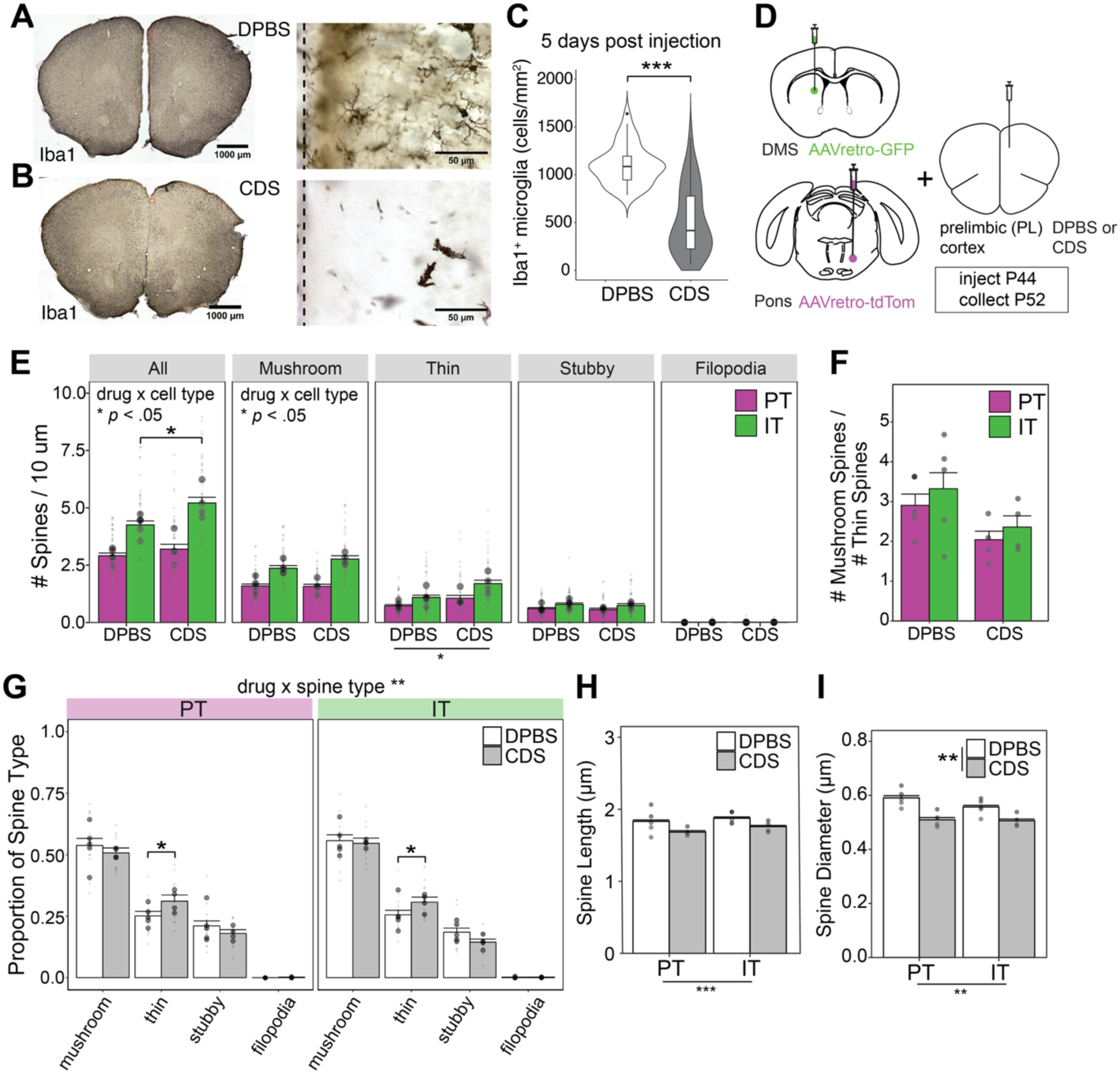
Effects of microglia depletion within prelimbic cortex on IT vs. PT spine density and morphology. Representative overview images of **A:** DPBS and **B:** CDS injected brain sections. **C:** Iba1+ cell density within the PL in DPBS vs. CDS injected mice, 5 days post-injection. **D:** Schematic of viral and drug infusions. **E:** Spine density for all spine types and for each class of spines. **F:** Ratio of mushroom to thin spines. Bar and error bars represent mean ± SEM of ratio for all dendrites. **G:** Proportion of each spine type was calculated as described in Fig. 2. **H:** Spine length and **I:** Spine diameter. For H and I, points represent mean values for each mouse. Bar and error bars represent mean ± SEM, and small points represent individual dendrites and larger points represent means for each mouse. For D-I: n = 4-5 mice / group, 24-30 brain sections / mouse, 48-60 dendrites / group, and > 900 spines / group. For all panels, * indicates *p* < .05, ** indicates *p* < .01, and *** indicates *p* < .001 for main effects, interaction effects, and *post hoc* comparisons from linear mixed effects models. See supplemental statistics file for full details.

For total spine density (Fig. 3E), we observed a significant main effect of cell type (*p* < .0001). Consistent with our previous experiment (Fig. 2E), spine density was higher on IT_DMS_ compared to PT_pons_ neurons (DPBS: *p* < .0001, CDS: *p* < .0001). In addition, there was a significant interaction between cell type and drug (*p* = .0245), whereby CDS injection was associated with higher total spine density in IT_DMS_ (*p* = .0143) but not PT_pons_ neurons (*p* = .40).

Spine type analysis indicated a significant interaction between drug and cell type for mushroom spine density (*p* = .0423). Mushroom spine density trended higher in CDS vs. DPBS injected mice on IT_DMS_ cells but the *post hoc* comparison was not significant (*p* = .0642) (Fig. 3E). This trend was absent for PT_pons_ neurons (*p* = .9314). Thin spine density showed significant main effects of drug (*p* = .0333) and cell type (*p* < .0001) but no interaction (*p* = .1141) (Fig. 3E). Stubby spine density showed only a main effect of cell type (*p* = .0026), with IT_DMS_ neurons again having a higher density than PT_pons_ neurons (Fig. 3E). No significant effects or interactions were detected in filopodia spine density. These data replicate IT_DMS_ vs. PT_pons_ neuron spine type density differences reported in Fig. 2E. Finally, analysis of the ratio of mushroom to thin spines (Fig. 3F) did not show significant main effects or interactions.

We next investigated how microglia ablation affected the proportion of each spine type and their morphological parameters. The mixed effects model for the proportion of each spine (Fig. 3G) type showed a significant interaction between spine type and drug (*p* = .0030). *Post hoc* comparisons between CDS and DPBS conditions within each spine type showed that CDS injection significantly increased the proportion of thin spines compared to DPBS injection (Fig. 3G). Since we observed cell type differences in the effect of CDS on total spine density, we performed *post hoc* comparisons between drug injections within each cell type and within each spine type. Whereas CDS increased total spine density only in IT_DMS_ cells, we observed CDS increased the proportion of thin spines in both cell types (PT_pons_: *p* = .0213; IT_DMS_: *p* = .0476) (Fig. 3G). This finding suggests that microglia may preferentially prune thin spines during the mid-adolescent to young adult transition and that microglia ablation during this window results in the persistence of thin spines.

Analysis of spine back bone length (Fig. 3H) showed a significant main effect of cell type only (*p* = .0005). Consistent with the findings of our developmental study (Fig. 2I), dendrites from IT_DMS_ cells had longer spines than spines from PT_pons_ cells (Fig. 3H). Although non-significant, there was a trend for CDS to decrease spine length (*p* = 0.05). The model for spine head diameter showed significant main effects of both drug injection (*p* = .0058) and cell type (*p* = .0040), and the interaction between these effects was approaching significance (*p* = .05). *Post hoc* comparisons showed that CDS injection significantly decreased spine head size in both cell types (IT_DMS_: *p* =.0177; PT_pons_: *p* = .0014) (Fig. 3I). *Post hoc* comparisons between cell types within each drug injection showed that in DPBS injected animals only, spines from PT_pons_ neurons have a larger head diameter (DPBS: *p* = .0005, CDS: *p* = .52). Overall, these data suggest that CDS decreased the length and size of dendritic spines in a non cell type-specific manner (Fig. 3H-I).

### 2.4: Role of mPFC microglia during adolescence in cognitive flexibility

Synaptic refinement within the mPFC during adolescence is believed to contribute to the maturation of executive function, including cognitive flexibility (Pöpplau et al. 2023). Given the observed effects of mPFC microglia ablation on dendritic spines, we next asked whether the same manipulation affected cognitive flexibility in the short or long term. Mice were bilaterally infused with 200 nL CDS or DPBS into the prelimbic cortex at P44 and then trained in an odor-based reversal learning task either 8 days later as adolescents (P52) or 6 weeks later as adults (P86) (Fig. 4A). We did not find evidence that mPFC microglia ablation affected performance during the discrimination or reversal phases of the task in terms of trials to criterion or proportion of errors (Fig. 4B-C), whether mice were trained shortly after microglia ablation (P52) or later in adulthood (P86) (3way ANOVA time x drug x sex, *p*’s > 0.05 all main effects and interactions; see Supplementary Statistics file for full details). Therefore, while CDS significantly altered spine density on IT_DMS_ neurons and spine type proportion on both IT_DMS_ and PT_pons_ neurons, this was not associated with enduring impairments in cognitive flexibility as assessed by the odor-based reversal learning task.

**Fig. 4:**
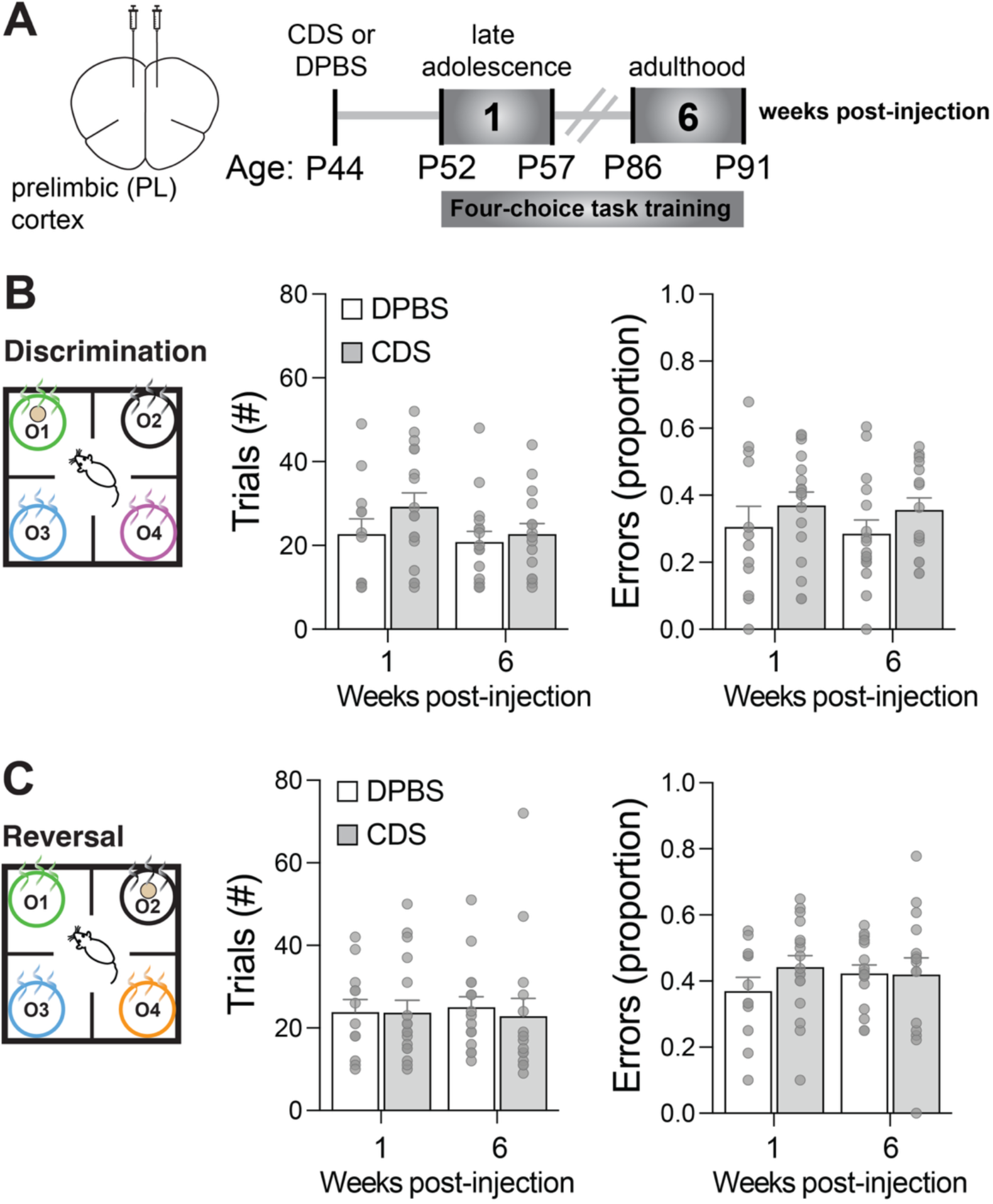
Transient prelimbic cortex microglia ablation during adolescence does not impair cognitive flexibility later in adolescence or adulthood. **A:** CDS or DPBS was bilaterally infused into PL at P44 and mice then underwent four-choice task training 1 week later form P52-57 or 6 weeks later from P86-91. **B:** Discrimination phase training. There was no main effect of drug, time, sex or interaction among factors on trials to criterion or proportion of errors (*p’s* > .05). **B:** Reversal phase training. There was no main effect of drug, time, sex or interaction among factors on trials to criterion or proportion of errors (*p’s* > .05). For B&C, n= 12-17 mice / group. See supplemental statistics file for full details.

## 3: DISCUSSION

Here we showed that IT- vs. PT-type pyramidal neurons exhibit distinct changes in dendritic spine properties across the adolescent transition and provide preliminary evidence that microglia may contribute to the cell type-specific pattern of spine maturation. Our study focused on apical dendritic spines, which *in vivo* imaging studies have demonstrated are associated with learning in motor cortex (Chen et al. 2015; Ma et al. 2016) and exhibit distinct plasticity rules compared to basal spines (Wright, Hedrick, and Komiyama 2025). In the context of the mPFC, studies have shown that apical dendritic spine plasticity on prelimbic cortex neurons is important for the consolidation of new memories relevant to goal-directed learning (Sequeira et al. 2023; Swanson, DePoy, and Gourley 2017). We observed that IT-type neuron spine properties were more dynamic during adolescence, showing significant changes in density, spine type proportion, and morphological parameters that were absent in PT-type neurons. A previous study found that Layer 3 and Layer 5 neurons in the mPFC showed similar developmental patterns in apical dendritic spine density during the early postnatal period (P6-30) (Kroon et al. 2019), suggesting that PT and IT cell types may diverge during adolescence, or that differences are only apparent at the level of spine types.

Dendritic spines can be classified into morphological categories (aka spine types) that when aligned with previous ultrastructural and *in vivo* imaging studies provides insight into their functional properties (Berry and Nedivi 2017). Thin spines are highly labile and hold the potential to transform into stable mushroom spines, thus representing a substrate for new synaptic connections and learning (Bourne and Harris 2007). We found that the ratio of mushroom:thin spines increased from P29 to P60 on both cell types, suggesting that synapses mature and become more stable over the adolescent transition. Notably, only IT-type neurons showed significant changes in the proportion of both mushroom (increase) and thin (decrease) spines across adolescence, reflecting greater structural reorganization compared to PT-type neurons. These results are consistent with *in vivo* 2-photon studies that have observed that the fraction of persistent vs. transient spines increases from weaning to adulthood in visual and somatosensory cortex (Holtmaat A et al. 2009; Grutzendler, Kasthuri, and Gan 2002; Trachtenberg et al. 2002; Zuo Y and Chang P 2005). These differing patterns of structural maturation suggest that IT-type neurons may retain a greater capacity for plasticity during adolescence. Chronic *in vivo* imaging studies, particularly during learning paradigms, are needed to further explore this hypothesis.

Our study adds to a growing body of data that implicates microglia in dendritic spine elimination within the frontal cortex during adolescence. Previous studies have observed that microglia exhibit less ramified morphology and enhanced phagocytic activity during adolescence (Mallya et al. 2019; Pöpplau et al. 2023). Mallya et al. observed that microglia within the mPFC preferentially engulfed longer, immature spines on Layer 5 pyramidal neurons (Mallya et al. 2019). Notably, we observed an increase in the proportion of thin spines at P52 when microglia were ablated at P44, suggesting that immature spines persisted that would have otherwise been eliminated. Whereas thin spine proportion increased on both IT_DMS_ and PT_pons_ neurons, only IT_DMS_ neurons showed a significant increase in total spine density (Fig. 3E), perhaps due to the baseline cell type differences in thin spine density. A previous study that examined the effect of microglia ablation within the mPFC during adolescence found that this led to a significant reduction in mushroom spine density on Layer 2/3 (IT-type) pyramidal neurons later in adulthood (Schalbetter et al. 2022). Our study focused on the short-term effect of microglia ablation, probing whether it affected the age-dependent decline we observed in thin spine density and thin spine proportion on IT-type neurons from mid-adolescence (P44) to late adolescence (P52). Future studies could examine how microglia ablation affects dendritic spine properties on IT vs. PT-type neurons later in adulthood. Our findings raise the question of what mechanisms lead could microglia to target thin, immature spines for elimination, and/or what mechanisms exist to promote spine loss on particular cell types. Potential mechanisms include differential expression of complement pathway proteins (Comer et al. 2020; Yilmaz et al. 2021), “eat-me” signals such as phosphatidylserine or complement component 3 (Scott-Hewitt et al. 2020; Schalbetter et al. 2022; Weinhard et al. 2018; Ball, Green-Fulgham, and Watkins 2022), or even microglia-specific receptor expression directing cell type-specific interactions (Favuzzi et al. 2021). Of note, synaptic engulfment does not appear to be required in order for microglia to influence dendritic spine elimination (Cheadle et al. 2020; Weinhard et al. 2018) and cell autonomous mechanisms, completely independent of glia, exist to promote spine elimination (Südhof 2018). Further studies are needed to understand the potential neuronal-glial interactions that promote cell type-specific spine elimination within the cortex during adolescence.

While we observed significant effects of microglia ablation on spine properties within mPFC, we did not observe behavioral effects in a cognitive flexibility task later in adolescence nor 6 weeks later in adulthood. The selected four-choice odor-based reversal learning task has previously been shown to depend on the mPFC and downstream DMS in juvenile and adult mice ( Johnson and Wilbrecht 2011; Delevich et al. 2022; Johnson et al. 2016). We selected this task because it is rapidly learned by mice, enabling behavioral assessment within a time window that is consistent with our observed spine changes following microglia ablation (∼8-10 days) (Fig. 3). Two recent studies reported impaired mPFC-dependent cognitive performance following transient microglia ablation during adolescence in mice. Schalbetter et al. found that CDS injection into mPFC at P42 led to deficits in mPFC-dependent tasks 6 weeks later in adulthood, including social recognition memory and temporal order memory (Schalbetter et al. 2022). Here the timing of microglia ablation and latency to behavioral testing were nearly identical to our 6 week cohort (Fig. 4). This suggests that while the mPFC has been implicated in the four-choice reversal learning task, microglia deficiency during mid-adolescence does not significantly impact performance in this task. Popplau et al. found that system administration of the colony-stimulating factor 1 receptor inhibitor, PLX3397, in early adolescence (∼P30) impaired late-adolescent (P56-60) performance in the same four-choice reversal learning task we examined (Pöpplau et al. 2023). While the authors observed accompanying changes in mPFC network activity, due to the brain-wide effect of PLX3397, it is possible that task impairment occurred downstream of microglia ablation effects within a different brain region. Alternatively, differences in age of manipulation and latency between microglia ablation and testing could account for the differences between Opatz et al. findings and ours.

There are several limitations to our study that warrant discussion. Given that we performed spine imaging in fixed tissue, our study is cross-sectional in nature and spine data represent only a snapshot in time. Therefore, our data do not provide insight into the spine dynamics – spine formation, elimination, and turnover – that contribute to the observed changes in density. Next, we used a retrograde AAV labeling approach with fluorescent nanobody enhancement to label dendritic spines in projection-defined cell populations. This labeling strategy does not label dendrites in a “Golgi-like” manner in the way that the Thy1-YFP-H transgenic line does (Feng et al. 2000). While the Thy1-YFP-H line brightly and sparsely labels PT-type pyramidal neurons, a similar transgenic reporter line is not currently available for IT-type neurons. Therefore, to achieve comparable labeling of IT and PT-type neurons, we used retrograde AAVs targeted to IT- and PT-specific projection targets. It is therefore possible that spines, particularly thin ones, may have gone undetected in the current study. However, we replicated previously reported differences in spine density measured by comparing Thy1-YFP-H+ (PT-type) to neurons *in utero* electroporated with fluorescent reporter constructs at E15.5 (IT-type) (Tjia et al. 2017) or when comparing biocytin-labeled neurons within mPFC between Layer 3 and Layer 5 (Kroon et al. 2019), suggesting that our labeling strategy was sensitive enough to detect these cell type differences. While the neuronal populations we imaged represent IT- and PT-type neurons, it should be noted that our results may not apply broadly to all IT- and PT-type neurons. Retrograde labeling and single axon tracing studies indicate that PT_pons_ neurons are part of the broader subclass of Layer 5b PT neurons that do not collateralize within basal ganglia (Economo et al. 2018). *In vivo* recordings from this tagged neuronal population suggest that PT_pons_ neurons are involved in later motor preparation and motor commands, although that study was performed in premotor cortex (Economo et al. 2018). Likewise, cross-corticostriatal (IT_DMS_) neurons represent a subclass of IT-type neurons. Retrograde AAV injection into contralateral DMS labeled IT-type neurons in Layer 2/3 as well as 5, and we did not distinguish soma location in our dendritic spine imaging experiments. One study found that in the secondary motor cortex, Layer 5 IT-type neurons were more likely to send axonal collaterals to contralateral striatum compared to Layer 2/3 IT-type neurons (Morita, Im, and Kawaguchi 2019), suggesting that our spine imaging results may represent more of the former population. However, we did not observe the corresponding differences in the laminar distribution of retrogradely-labeled IT_DMS_ neurons – *i.e.* cell body positions were not deeper in the contralateral compared to ipsilateral hemisphere. Here, we manipulated microglia and then visualized dendritic spine changes, but future studies should more directly examine microglia internalization of labeled IT vs. PT-type neuron material (Mallya et al. 2019; Comer et al. 2020; Yilmaz et al. 2021; Pöpplau et al. 2023).

The current study builds on our previous work examining the influence of gonadal hormones at puberty on the adolescent pruning of apical dendritic spines on IT_DMS_ neurons in mPFC (Delevich, Okada, et al. 2020). In that study we employed the same retrograde AAV strategy to label IT_DMS_ and PT_pons_ neurons, verifying with slice electrophysiology that the labeled pyramidal neuron populations displayed their subtype’s characteristic membrane properties. There, our spine analysis was restricted to IT_DMS_ neurons, we examined only two timepoints (P29 vs. P60), and we did not assess spine morphological parameters. There were two findings from Delevich, Okada et al. that we did not replicate here: first, we did not find a main effect of sex on dendritic spine density on IT_DMS_ neurons, and second, we did not find a significant decrease in *total* spine density from P29 to P60. Our current findings are in line with what was previously reported in a study examining apical spine density of Golgi-impregnated Layer 2/3 pyramidal neurons in mPFC of rats across adolescence: spine density significantly increased between P20 and P30, but did not subsequently change at P56 or P90 (Markham, Mullins, and Koenig 2013). Another study observed that puberty onset was associated with a significant decrease in the number of synapses labeled by synaptophysin in the mPFC of male and female rats (Drzewiecki CM 2016). Here we found that mushroom spine density on IT_DMS_ neurons increased from P29 to P44 while thin spine density decreased by P60 compared to P29 and P44. While we did not assess fully assess pubertal development, we confirmed in a separate group of mice that no evidence of mature spermatids (Spears, Matthews, and Hartwig 2013) or corpus lutea (Gaytan et al. 2017) were seen at P29 in male and female mice, respectively. Therefore, this work points to mid-adolescence, when pubertal maturation occurs, as an inflection point when spine density ceases to increase. Our previous study found that prepubertal gonadectomy in male mice resulted in higher apical spine density on IT_DMS_ neurons in young adulthood, suggesting that puberty plays a role in spine maturation, at least in males (Delevich, Okada, et al. 2020). Next, there is an important methodological difference between studies: we previously performed imaging on a 2-photon microscope (Delevich, Okada, et al. 2020), whereas here we used a confocal microscope with image deconvolution. We previously reported slightly higher spine densities, perhaps due to the better spatial resolution afforded by 2-photon. In the current study, we observed more subtle sex differences, whereby females tended to have a higher proportion of thin spines relative to mushroom and exhibited greater cell type differences in spine density. If our current labeling and imaging strategy was more limited in detecting dimmer, thinner spines, this could have disproportionately affected females, thus minimizing sex differences in overall spine density.

To the best of our knowledge, our study is the first to directly compare cortical IT vs. PT type pyramidal neuron spine properties across the adolescent transition. Our study adds to a growing body of research demonstrating that spine density and dynamics change within agranular frontal cortices of mice during adolescence (Zuo Y and Chang P 2005; Gourley et al. 2012; Johnson, Peckler, and Janak PH 2016; Pattwell et al. 2016; Boivin et al. 2018; Pöpplau et al. 2023) and provides new insight into pyramidal cell type-specific spine maturation. Given that we observed frontal cortical IT-type neurons undergo greater changes in apical dendritic spine properties during adolescence compared to PT-type neurons, this suggests they may be more vulnerable to enduring effects of endogenous or exogenous influences during adolescence, for instance genetic factors vs. drugs or stressors. Due to the deep conservation of the broad pyramidal cell types investigated (Hodge et al. 2019), our data may also provide insight into the cellular processes that underlie cortical maturation in humans.

## Supporting information

Supplementary Statistics

## Acknowledgments

We thank Alexis Daniels and Theodora Gill for technical assistance. We thank Dr. Sara Westbrook for feedback on the manuscript.

## CRediT statement

Klappenbach (Investigation, Data Curation, Formal analysis, Visualization, Writing – original draft), Wang (Investigation), Lenkersdorfer (Investigation), Buursma (Methodology), Acuna (Investigation), Chu (Data curation, Investigation), Chen (Formal Analysis), Richards (Investigation), Herrera (Investigation), Touretsky (Investigation), and Delevich (Conceptualization, Funding Acquisition, Supervision, Visualization, Writing – original draft).

