## Supplementary Materials for "Cell Type-Specific Changes in Dendritic Spines Across Adolescence Within Mouse Medial Prefrontal Cortex"

Supplementary Methods:

### Animals

Experiment 1 examined the spatial distribution of IT and PT neurons and consisted of 2 females and 2 males aged P60-100 at the time of perfusion. Experiment 2 examined spine density on IT and PT neurons across adolescence and consisted of the following mice: 5 P29 females, 6 P29 males, 4 P44 females, 5 P44 males, 5 P60 females, and 6 P60 males. Experiment 3 quantified CDS ablation of microglia at 5 days post-injection and consisted of 6 CDS injected mice and 7 DPBS injected mice. Experiment 4 determined the effects of CDS-mediated microglia ablation on IT and PT neuron spine density and consisted of 4 CDS injected mice and 5 DPBS injected mice balanced by sex. Finally, Experiment 5 assessed the effects of CDS-mediated microglia ablation of cognitive flexibility assessed by the four-choice odor-based reversal learning task. It consisted of 12 adolescent DPBS and 17 adolescent CDS mice and 16 adult DPBS and 15 adult CDS mice.

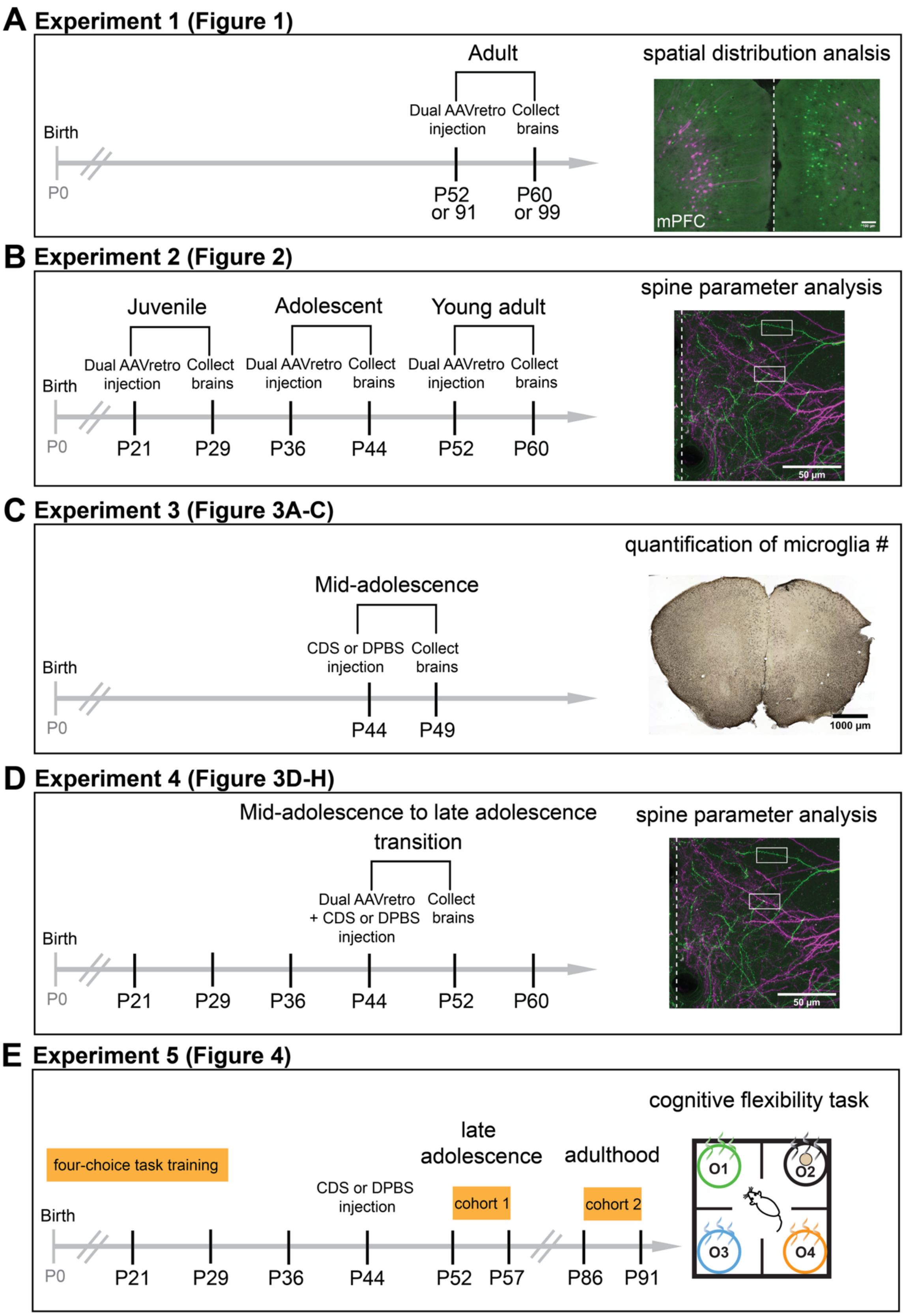


**Supplemental Fig. 1:** Timeline of all experiments performed.

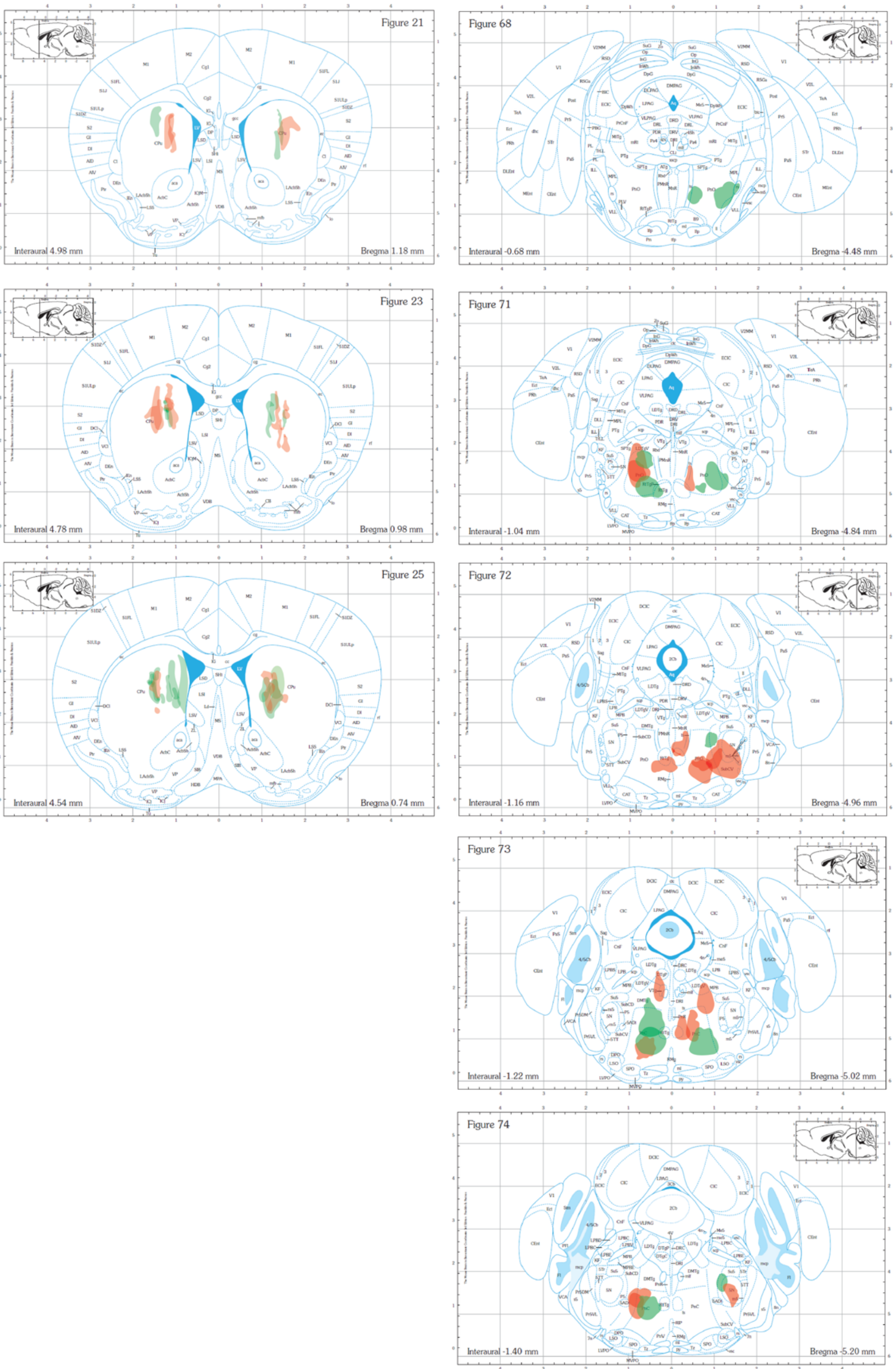


**Supplemental Fig. 2: Viral vector spread for developmental dendritic spine imaging experiments.** Viral vector spread maps for mice in main text Figure 2, with DMS injection sites shown in the left column and Pons injection sites shown in the right column. Color reflects fluorophore injected, and the opacity of all viral spread traces were set to 30% for visibility.

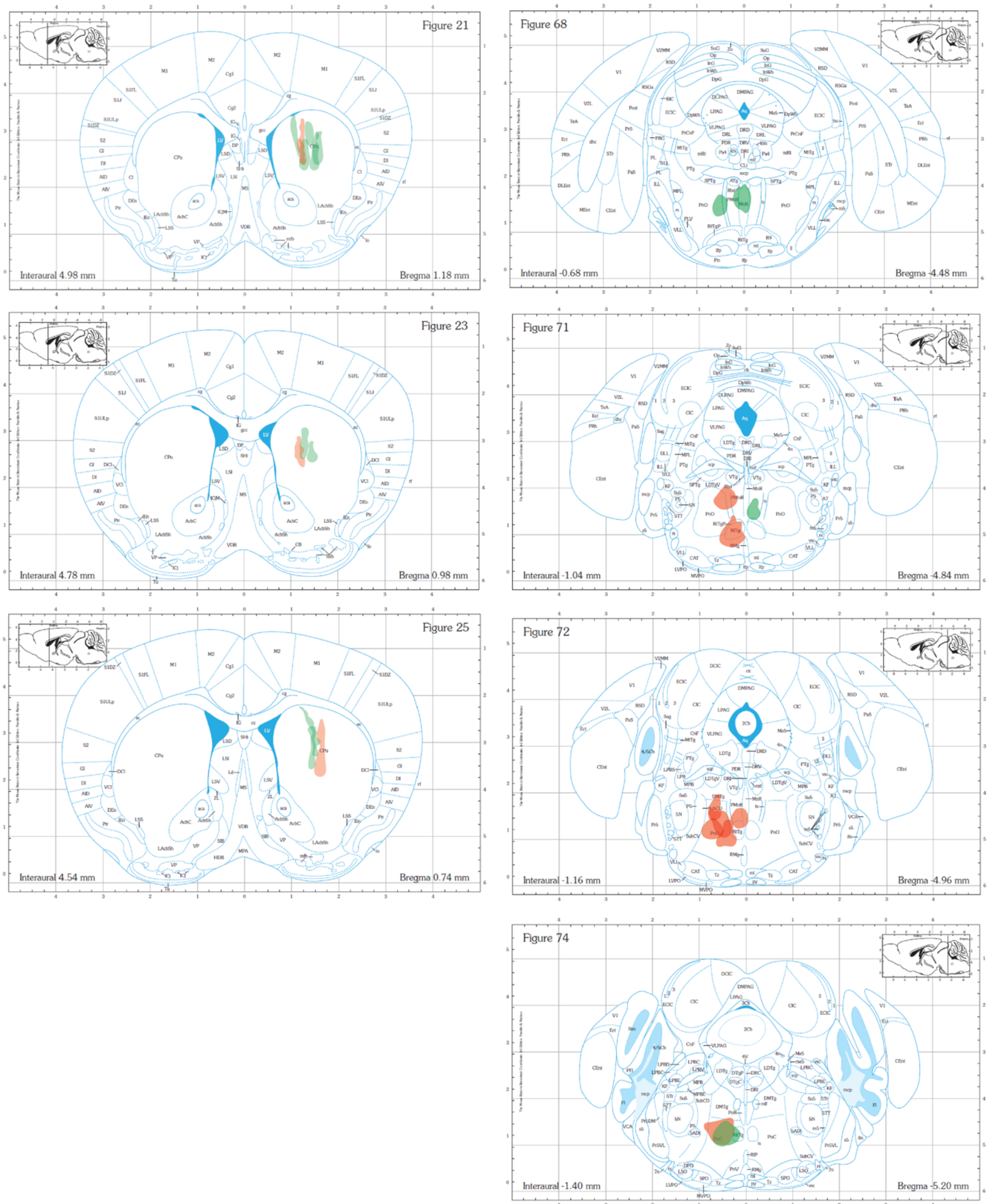


**Supplemental Fig. 3: Viral vector spread for microglia ablation with dendritic spine imaging experiments.** Viral vector spread maps for mice in main text Figure 3, with DMS injection sites shown in the left column and Pons injection sites shown in the right column. Color reflects fluorophore injected, and the opacity of all viral spread traces were set to 30% for visibility.

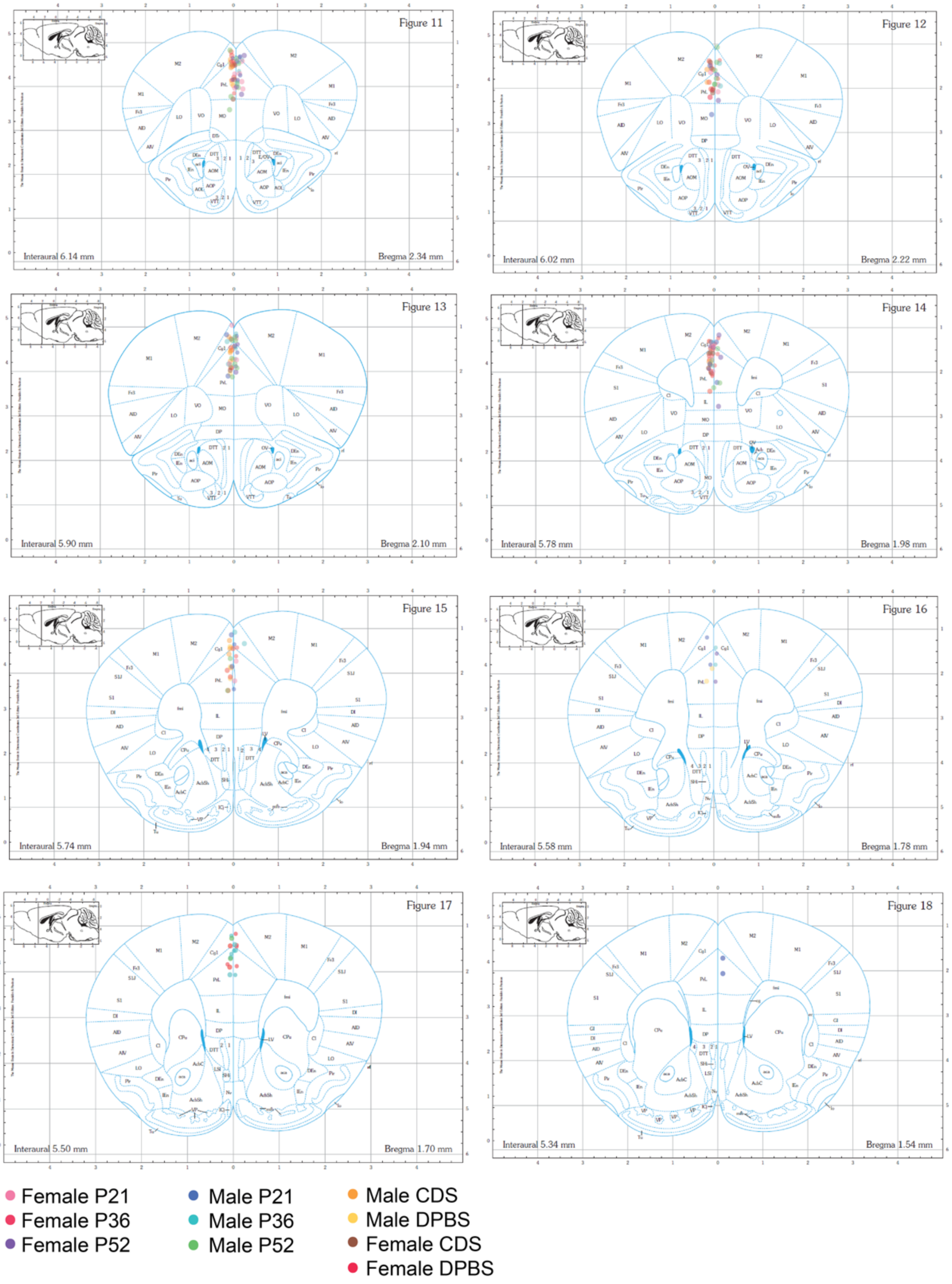


**Supplemental Fig. 4: Confocal imaging sites for dendritic spine analysis.** Related to data presented in Figures 2 & 3. A bleach spot was created in a distal location to each region of interest (ROI) by scanning the 405 nm laser at 100% power for 1-2 minutes. The coordinates of each ROI and bleach spot were recorded and then used to map the imaging locations to the Paxinos brain atlas. Colored dots correspond to sex, age, and in cases related Figure 3, mPFC drug injection.

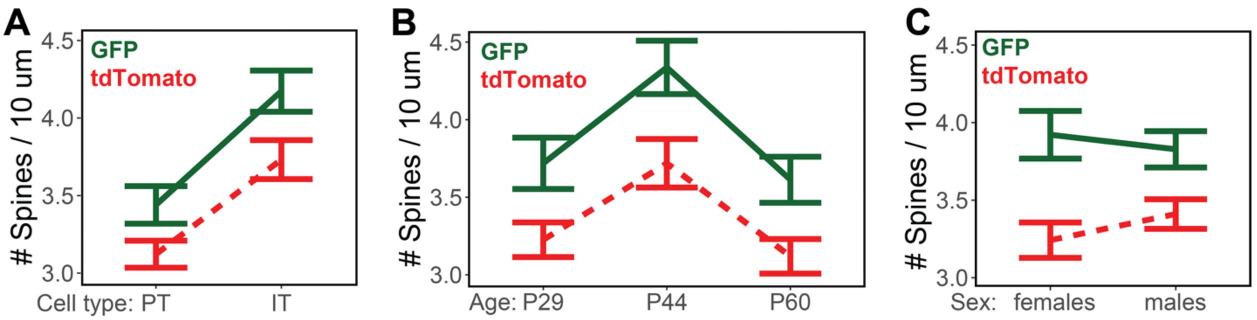


**Supplemental Fig. 5:** Effect of flurophore color on spine density. Plots show effect of flurorphore color on other model effects (**A:** cell type, **B:** age, **C:** sex) collapsed across other variables. GFP is shown with solid line and tdTomato is shown with dashed line. Lines and error bars represent mean ± SEM for all dendrites. n = 9-19 mice / group, > 100 dendrites / group.

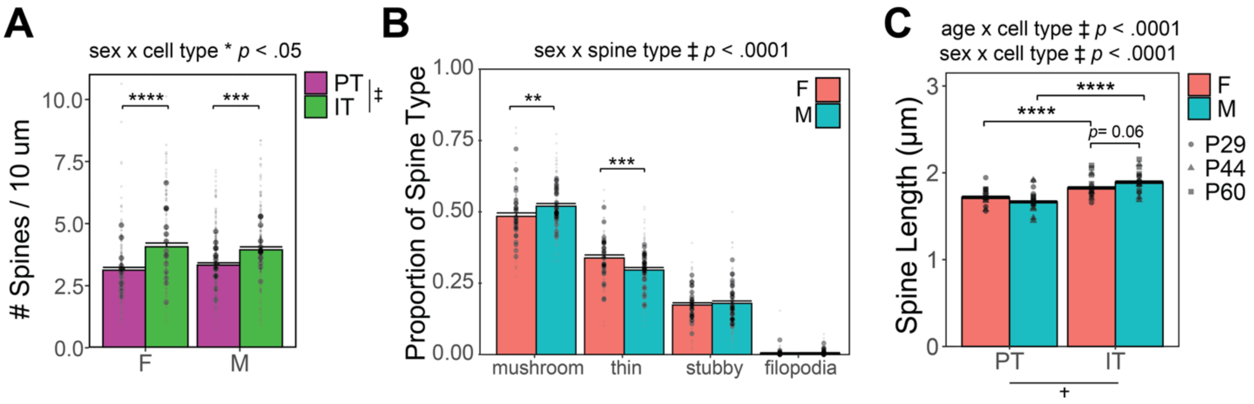


**Supplemental Fig. 6:** Sex differences in spine parameters. **A:** Spine density for all spine types. Small points represent individual dendrites and larger points represent means for each mouse. Bar and error bars represent mean ± SEM of dendrites. **B:** Proportion of each class of spine type. Points represent means for each mouse and bar and error bars represent mean ± SEM of proportions for all brain sections. **C:** Spine length. Points represent mean for each mose and bar and error bars represent mean ± SEM of all spines. n = 14-17 mice / group, 40-50 brain sections, > 150 dendrites / group, > 2300 spines / group. For all panels, * indicates $p < .05$, ** indicates $p < .01$, *** indicates $p < .001$, and ‡ indicates $p < .0001$ for main effects, interaction effects, and *post hoc* comparisons from linear mixed effects models. See main text and supplemental statistical file for more details on statistical analysis and values.
